# Identification and neuroprotective properties of NA-184, a selective calpain-2 inhibitor

**DOI:** 10.1101/2023.10.16.562509

**Authors:** Michel Baudry, Yubin Wang, Xiaoning Bi, Yun Lyna Luo, Zhijun Wang, Zeechan Kamal, Alexander Shirokov, Ed Sullivan, Dennis Lagasca, Hany Khalil, Gary Lee, Kathy Fosnaugh, Philippe Bey, Shujaath Medi, Greg Coulter

**Author notes:** **Correspondence:** Michel Baudry CDM, Western University of Health Sciences 309 E 2^nd^ St Pomona, CA 91766.

## Abstract

Our laboratory has shown that calpain-2 activation in the brain following acute injury is directly related to neuronal damage and the long-term functional consequences of the injury, while calpain-1 activation is generally neuroprotective and calpain-1 deletion exacerbates neuronal injury. We have also shown that a relatively selective calpain-2 inhibitor, referred to as C2I, enhanced long-term potentiation and learning and memory, and provided neuroprotection in the controlled cortical impact (CCI) model of traumatic brain injury (TBI) in mice. Similar results were also obtained in conditional calpain-2 knock-out mice. Using molecular dynamic simulation and Site Identification by Ligand Competitive Saturation (SILCS) software, we report here the discovery of a selective calpain-2 inhibitor NA-184, (S)-2-(3-benzylureido)-N-((R,S)-1-((3-chloro-2-methoxybenzyl)amino)-1,2-dioxopentan-3-yl)-4-methylpentanamide whose properties make it a good clinical candidate for the treatment of TBI.

## Introduction

TBI is a significant public health problem in the United States. There were over 220,000 TBI-related hospitalizations in 2019 and 69,473 TBI-related deaths in 2021, making it the 8^th^ largest cause of death (https://www.cdc.gov). Among the different types of TBI, penetrating traumatic brain injuries produce the worst outcomes and highest mortality rates (Binder et al., 2005). TBI induces immediate neuropathological consequences, including neurodegeneration (Gupta and Sen, 2016) and axonal damage (Siedler et al., 2014).

Numerous reviews have discussed the role of calpain in neurodegeneration in general (Vosler et al., 2009) (Yildiz-Unal et al., 2015) in general, in stroke (Koumura et al., 2008; Anagli et al., 2009), and TBI (Liu et al., 2014; Kobeissy et al., 2015). Consequently, numerous studies have attempted to reduce neurodegeneration in both stroke and TBI using calpain inhibitors (Cagmat et al., 2015; Siklos et al., 2015). Results of these studies have been ambiguous, as some studies using the first generation of calpain inhibitors in TBI reported beneficial effects (Thompson et al., 2010), while other studies did not (Bains et al., 2013; Schoch et al., 2013). Importantly, all these studies used non-selective calpain inhibitors, which did not differentiate calpain-1 and calpain-2, the major calpain isoforms in the brain. Several reasons could account for the failure to develop clinical applications with such inhibitors, including their lack of specificity/potency/selectivity (Donkor, 2011), and the incomplete knowledge regarding the functions of calpain-1 and calpain-2 (aka µ- and m-calpain) in the brain. Work from our laboratory over the last 10 years has revealed that calpain-1 and calpain-2 play opposite roles in both synaptic plasticity and neuroprotection/neurodegeneration (Baudry and Bi, 2016). Thus, calpain-1 activation is required for theta burst stimulation-induced long-term potentiation (LTP) and is neuroprotective (Wang et al., 2013; Wang et al., 2014). On the other hand, calpain-2 activation limits the magnitude of LTP and is neurodegenerative (Wang et al., 2013; Wang et al., 2014). These findings could explain the failure of the previous studies to convincingly demonstrate the role of calpain in neurodegeneration and the lack of clear efficacy of the previously tested calpain inhibitors, which did not discriminate between calpain-1 and calpain-2.

We found that the peptidyl α-ketoamide, Z-Leu-Abu-CONH-CH_2_-C_6_H_3_(3,5-(OMe)_2_) (Li et al., 1996), referred to as C18 (called C2I above and shown in Fig. 1), with a reported 100-fold difference in Ki values for calpain-2 vs calpain-1-, did enhance the magnitude of TBS-induced LTP (Wang et al., 2014) and facilitated learning and memory (Liu et al., 2016). It also provided significant neuroprotection in both a mouse model of acute glaucoma (Wang et al., 2016) and a mouse model of TBI (Wang et al., 2018). However, in all our experiments, this compound exhibited an inverted U-shape dose response curve, with low doses providing learning facilitation and neuroprotection and high doses eliciting the opposite effects, which reflected the inhibition of calpain-2 and calpain-1, respectively. Based on this finding we initiated a program of compound optimization to identify a selective calpain-2 inhibitor with better properties for further drug development for the treatment of TBI.

**Figure 1.**
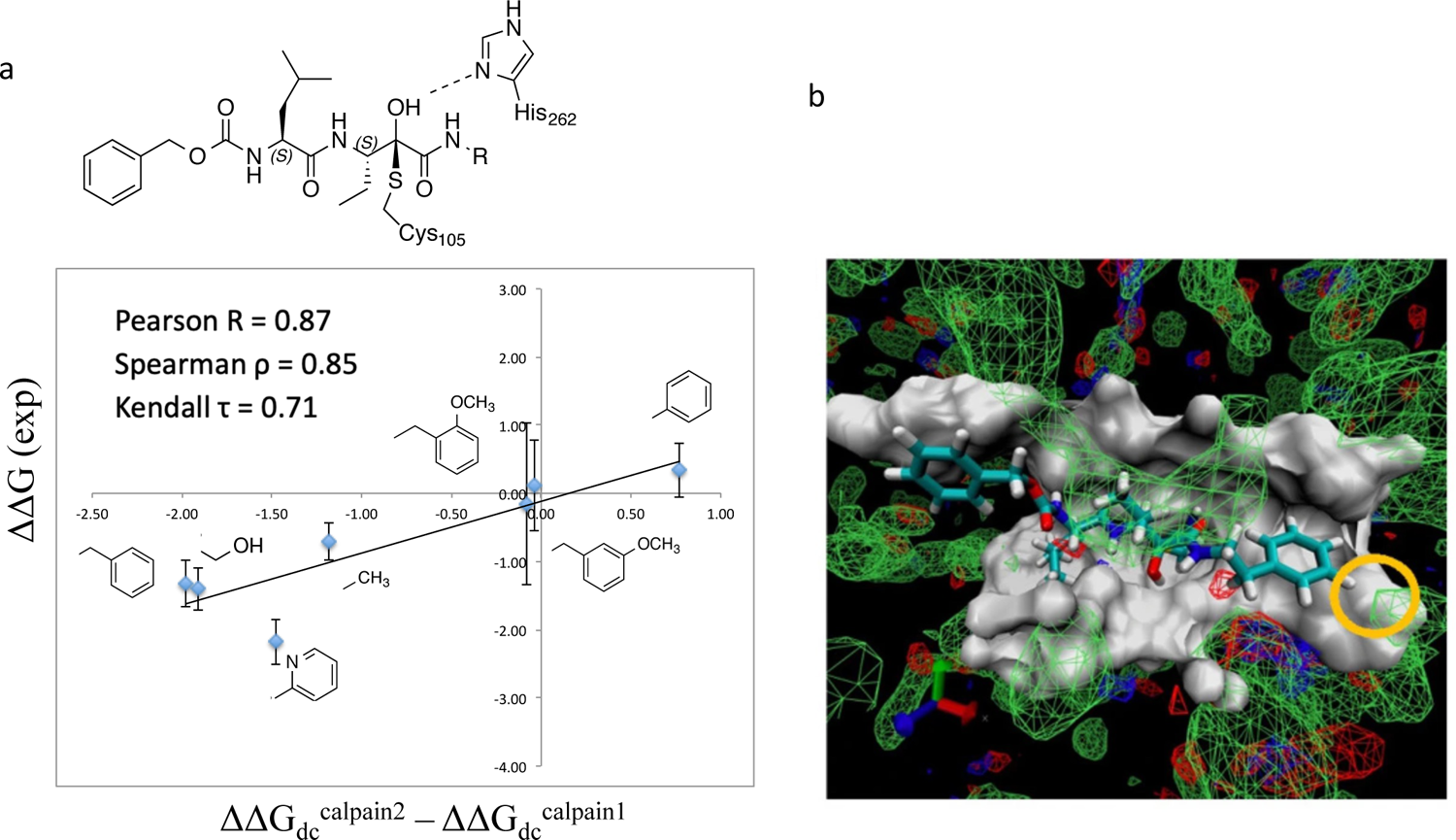
Molecular modeling of ketoamide calpain inhibitors. **a.** Chemical structures of the common core bound to the catalytic residues in calpain-2. Experimental selectivity data were obtained using ΔΔG (exp) = RT ln(*K*_i_^Calpain2^/ *K*_i_^Calpain1^), in which *K*_i_^Calpain2^ and *K*_i_^Calpain1^ are the inhibitory constants for each compound in calpain-2 and calpain-1 from Li et al., 1996. Calculated selectivity data were obtained using ΔΔG^calpain2^ – ΔΔG^calpain1^, in which ΔΔGs are the relative binding free energy difference between the core structure and each ligand in calpain-1 or calpain-2, attained via the FEP calculations. All free energy units are kcal/mol. Figure reproduced from Chatterjee, Botello-Smith et al. 2017. b. SILCS FragMaps of calpain-2. Green mesh density is polar FragMap, blue is hydrogen bond donor FragMap, and red is hydrogen bond acceptor FragMap. The cutoff value for all density mesh is set to be − 1. Yellow circles indicate para-position at P1’.

## Materials and Methods

### Molecular dynamics simulation

The crystal structures of rat calpain-1 (PDB 2R9C) and rat calpain-2 (PDB 3BOW) are mutated *in silico* to human’s using *Molecular Operating Environment* (MOE) (Molecular Operating Environment (MOE) v. 2013.08 (Chemical Computing Group Inc., 2016) to be consistent with the experimental assay. CHARMM36 force field (Best et al., 2012) was used for all simulations. CHARMM36 force fields for deprotonated cysteine and protonated histidine were used for the catalytic site (Chatterjee et al., 2017). The ketoamide warhead was re-parameterized previously (Chatterjee et al., 2017). All the MD simulation systems in explicit solvent were prepared by using the CHARMM-GUI (Jo et al., 2008). Each system was solvated into a rectangular water box consisted of CHARMM TIP3P water molecules (Jorgensen et al., 1983) and 150 mM KCl, with an edge distance of 10 Å. All the MD simulations were performed using NAMD (Phillips et al., 2005) under periodic boundary conditions at a constant temperature of 300 K and pressure of 1 atm (NPT ensemble) (Adelman and Garrison, 1976). A smoothing function was applied to van der Waals forces between 10 and 12 Å. The solvated complexes were minimized and equilibrated using a stepwise procedure set up by CHARMM-GUI.

To assist our medicinal chemistry campaign, we used Site Identification by Ligand Competitive Saturation (SILCS) program (Hess et al., 2008) to rank and prioritize chemical modification. Our previous study showed that docking the P1’ fragments produced higher ranking accuracy among reversible covalent analogs than docking the whole ligand in the calpain catalytic site [PMID: 30763080]. Hence, the binding affinity ranking was done following the protocol described in ref [PMID: 30763080].

### Analog synthesis

Synthesis of analogs of C2I was performed using traditional methods for peptidomimetics. All materials were obtained from commercial sources. The purity of each compound was checked by HPLC and ^1^H NMR and mass spectrometry. We are reporting below the synthesis of two of these analogs, which represent the most extensively studied members of the family of molecules synthesized in this project.

#### 1. Preparation of NA-112

Compound NA-112 was prepared in 10 steps from commercially available raw materials as summarized in Scheme 1.

**Scheme 1.**
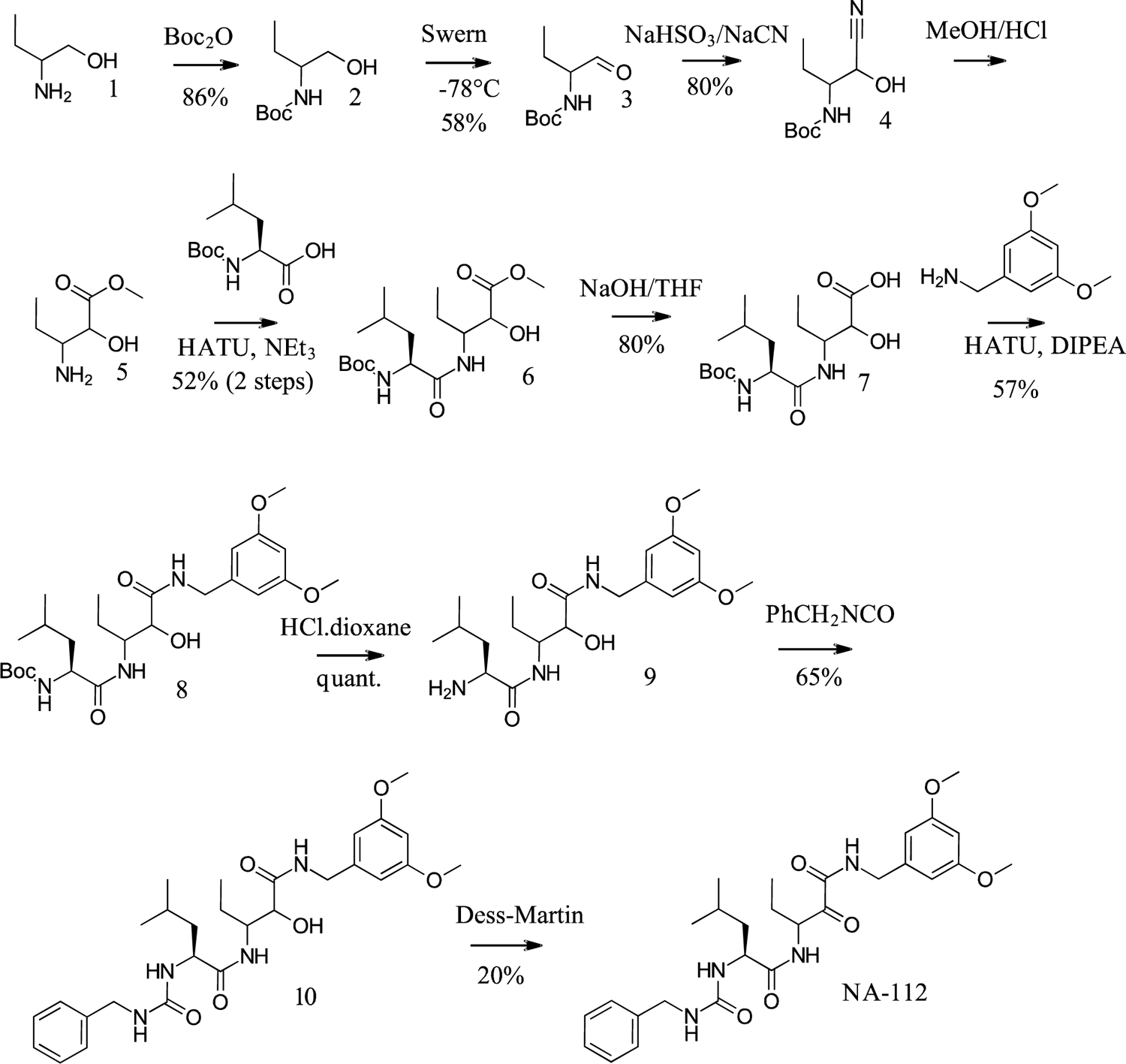

The final product was purified by column chromatography on silica gel, followed by recrystallization from dichloromethane to provide an off-white solid (98.4% purity by LC/MS). The product exists as a mixture of diastereomers, with the (S)-Leu center fixed.

#### 2. Preparation of NA-184

Compound NA-184 was prepared in 11 steps from commercially available raw materials as summarized in Scheme 2.

**Scheme 2.**
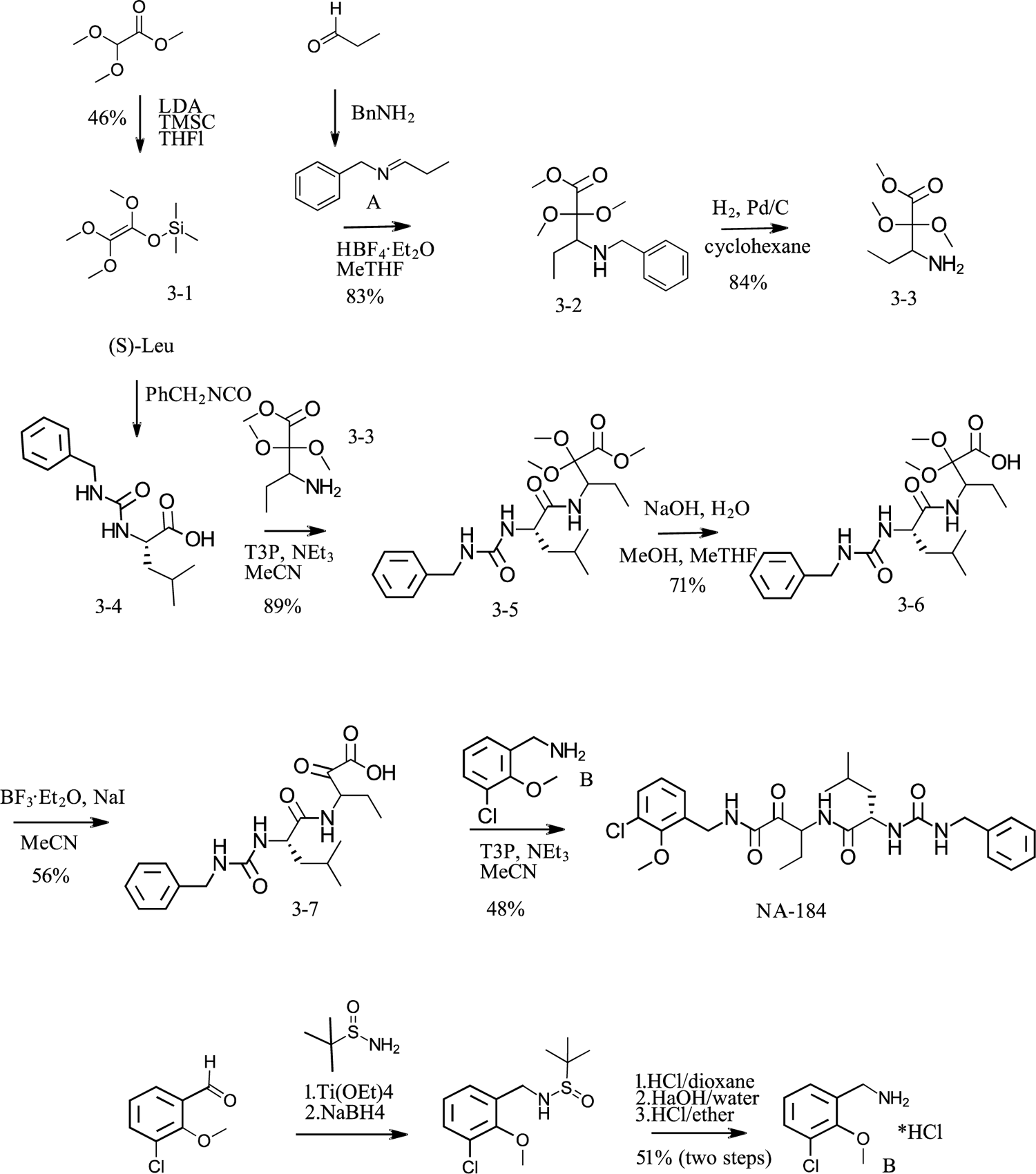

The final product was purified by column chromatography on silica gel, followed by precipitation from a mixture of MeTHF and MTBE to provide an off-white solid (99.1% purity by HPLC). The product exists as a mixture of diastereomers, with the (S)-Leu center fixed.

### Calpain assays

– *In vitro calpain assays:* The hydrolysis of the fluorogenic substrate Suc-Leu-Tyr-AMC by calpain-1 and calpain-2 was performed as previously described (Sasaki et al., 1984), with minor modifications. Briefly, purified calpain-1 from human or porcine erythrocytes (8 µg, Millipore) or human or rat recombinant calpain-2 produced by bacterial expression as described by (Elce et al., 1995) (plasmids were generous gifts from Dr. Peter Davies from Queen’s University, Ontario, CA) was incubated with Suc-Leu-Tyr-AMC (0.5 mM) in 60 mM imidazole-HCl buffer, pH 7.3, containing 5 mM CaCl_2_, 5 mM cysteine, 2.5 mM β-mercaptoethanol, and different concentrations the various molecules (ranging from 0 to 20 µM). The reaction was initiated by adding the enzyme and continued at 30 °C for 15 min, while the fluorescence of 7-amino-4-methylcoumarin (Ex 380 nm/Em 450 nm) was monitored every 30 sec in a POLARstar Omega fluorescence microplate reader (BMG Labtech). The rate of hydrolysis (increase in fluorescence/min) was determined from the linear portion of the curve. The IC_50_ values were obtained by adjusting data from each experiment into a sigmoidal dose-response curve. The Ki values were calculated from the average of the IC_50_ values and from a single substrate concentration by using a K_i_ calculator tool for fluorescence-based competitive binding assays (http://sw16.im.med.umich.edu/software/calc_ki/). The Km values of Suc-Leu-Tyr-AMC used for the Ki calculation were 4.74 mM and 2.21 mM for calpain-1 and calpain-2, respectively, as previously reported (Sasaki et al., 1984).
– *Ex-vivo calpain assays:* Mouse pons-cerebellum was used to provide a source of endogenous calpain-1 and calpain-2, as these regions exhibit the highest levels of calpain activity (Simonson et al., 1985). Membrane fractions were prepared by centrifugation of homogenates and were resuspended in 60 mM imidazole-HCl buffer, pH 7.3, containing 5 mM cysteine, 2.5 mM β-mercaptoethanol. Calpain-1 activity was measured by adding 20 µM calcium and the fluorogenic substrate (Suc-Leu-Tyr-AMC, 0.5 mM), while total calpain (calpain-1 and calpain-2) was measured in the presence of 5 mM calcium. Calpain-2 activity was calculated from the difference in activity between total calpain and calpain-1 activity. Alternatively, membrane fractions were prepared from pons-cerebellum from calpain-1 knock-out mice to measure calpain-2 activity.
– *Calpain assay using human cells:* We used HEK cells lacking either calpain-1 or calpain-2 (generous gift from Dr. Sandra Cooper, University of Sydney, Australia) to test the efficacy of NA-112 and NA-184 to inhibit calpain-mediated spectrin truncation. Cells were homogenized in calpain assay buffer and incubated in the presence of various concentrations of inhibitors and in the absence or presence of calcium (5 mM). Aliquots were processed for Western Blots with spectrin antibodies (1:1000, MAB1622, EMD Millipore).

### Various protease assays

Commercially available kits were purchased to test the effects of calpain inhibitors on various proteases.

### CCI model of TBI

#### Animals

Animal use in all experiments followed NIH guidelines and all protocols were approved by the Institution Animal Care and Use Committee of Western University of Health Sciences. Calpain-1 KO mice on a C57Bl/6 background were obtained from a breeding colony established from breeding pairs generously provided by Dr. Chishti (Tufts University). C57Bl/6 mice were purchased from Jackson Labs and were the corresponding wildtype (WT). Six-seven-month-old Sprague-Dawley rats (males and females) were also used for the controlled cortical impact (CCI) model of TBI.

#### Controlled cortical impact

CCI model was established in mice following the previously described protocol described (Wang et al., 2018). Mice (3-month-old, 25-30 g) or rats were anesthetized using isoflurane and fixed in a stereotaxic frame with a gas anesthesia mask. A heating pad was placed beneath the body to maintain body temperature around 33-35 °C. The head was positioned in the horizontal plane. The top of the skull was exposed, and a 5-mm craniotomy was made using a micro drill lateral to the sagittal suture and centered between Bregma and Lambda. The skull at the craniotomy site was carefully removed without damaging the dura. The exposed cortex was hit by a pneumatically controlled impactor device (AMS-201, Amscien). The impactor tip diameter was 3 mm, the impact velocity was 3 m/sec, and the depth of cortical deformation was 0.5 mm. After impact, the injured region was sutured using tissue adhesive (3M) and mice were placed in a 37 °*C* incubator until they recovered from anesthesia. In sham surgery, animals were sutured after craniotomy was performed. The selective calpain-2 inhibitors were injected in general intraperitoneally (ip) at various doses and control animals were injected with the indicated vehicles (5% DMSO in PBS).

### Immunohistochemistry (IHC)

– *Spectrin breakdown product:* Twenty-four h after TBI, mice or rats were anesthetized and perfused intracardially with freshly prepared 4% paraformaldehyde in 0.1 M phosphate buffer (PB, pH 7.4). After perfusion, brains were removed and immersed in 4% paraformaldehyde at 4 °C for post-fixation, then in 15% and 30% sucrose at 4 °C for cryoprotection. Three coronal frozen sections (20 µm thick) in each brain at Bregma −0.58, −1.58, −1.94 were prepared. In order to determine in situ calpain activation, sections were co-stained with rabbit anti-SBDP (1:500, a gift from Dr. Saido, Riken, Japan) antibodies. Sections were first blocked in 0.1M PBS containing 5% goat or donkey serum and 0.3% Triton X-100 (blocking solution) for 1 h, and then incubated with primary antibodies prepared in blocking solution overnight at 4 °C. Sections were washed 3 times in PBS (10 min each) and incubated in Alexa Fluor 488 goat anti-rabbit IgG, prepared in blocking solution for 2 h at room temperature. After three washes, sections were mounted with mounting medium containing DAPI (Vector Laboratories). For quantification of SBDP levels, three 637 µm x 637 µm areas near the lesion site in each section were imaged using confocal microscopes (Nikon C1 and Zeiss LSM-880) and analyzed. In each area, the regions proximal (0 - 170 µm from the lesion edge) and distal (> 170 µm from the lesion edge) to the impact site were separately outlined using the “freehand selection” function of ImageJ, and mean fluorescence intensity (MFI) of SBDP was measured. Data in all three sections from the same brain were averaged. For quantification of GFAP and iba1 levels, whole brain sections were imaged using the Tile Scan function of an LSM 510 confocal microscope (Zeiss). Three 300 µm x 300 µm areas centered 400 µm to the lesion edge on the ipsilateral side and three areas of the same size centered 700 µm to the cortical surface in the contralateral side were selected. MFI of GFAP or iba1 in each area was measured in ImageJ. Then, MFI in the contralateral side (background signal) was subtracted from MFI in the ipsilateral side. Data in all three sections from the same brain were averaged. Image acquisition and quantification were done by two persons in a blind fashion.
– *TUNEL staining.* Brains were collected at 0, 6, 24 and 72 h after TBI. Coronal frozen sections (20 µm thick) at Bregma 1.54, 0.50, −0.58, −1.58, −1.94 and −2.30 were prepared. TUNEL staining was performed in a set of sections using the ApopTag *in situ* apoptosis detection kit (S7165, Millipore). Sections were visualized under confocal microscopy (Nikon). All TUNEL positive nuclei surrounding the lesion area in 6 sections of each brain were counted using the “analyze particles” function in ImageJ. To separately analyze cell death in the region proximal and distal to the impact site, the regions proximal (0 - 170 µm from the edge of impact site) and distal (> 170 µm from the edge of impact site) to the impact site were outlined using the “freehand selection” function of ImageJ. TUNEL positive cells in these two regions were separately counted. Image acquisition and quantification were done by two examiners in a blind fashion.

### Pharmacokinetics and pharmacodynamics

The pharmacokinetic study of NA-184 was carried out in mice by intravenous injection via tail vein with a dose of 10 mg/kg in a liposomal formulation. Blood samples were collected before dosing (0 time) and at time points of 1, 5, 15, 30 min, 1, 2, 4, 8, 16, 24 hr post dose. Approximately, 0.8 ml of blood were collected into heparinized tubes via cardiac puncture. Plasma was separated by centrifuging for 5 min at 10,000 rpm. Whole brains were collectedwhich were weighed and mixed with 3x (by volume) PBS buffer to homogenize. All samples were kept at −80 °C before further processing.

The concentration of NA-184 was determined using an HPLC-MS/MS method with NA-101 as the internal standard. Briefly, NA-184 was extracted using a liquid-liquid extraction method. For each 100 μl of plasma samples or tissue homogenate, 10 μl of the internal standard solution (1000 ng/ml) was added. After mixing, 400 µl methyl tert-butyl ether was added to each sample mixture. The samples were then mixed using a vortex mixer for 3 min followed by centrifuging at 5,000 rpm for 3 min. The upper clear solvent layer was transferred into a clean glass tube and evaporated under the nitrogen blow until dry. Then 0.1 ml of 80% acetonitrile was added to each tube and vortexed for 1 min. The mixture was centrifuged at 10,000 rpm for 5 min and 5 µl injected into the HPLC-MS/MS system for analysis.

The LC/MS/MS system consisted of an API 3200 LC/MS/MS system (Sciex, Framingham, MA) and two Shimadzu LC-20AD Prominence Liquid Chromatograph pumps equipped with an SIL-20A Prominence autosampler (Shimadzu, Columbia, MD, USA). Chromatography was carried out using a Zorbax SB C18 column (150 × 2.1 mm, 5 µm, Zorbax, Agilent, Santa Clara, CA, USA) which was proceeded with a SB-C18 Guard Cartridges (12.5 × 2.1 mm, Zorbax, Agilent, Santa Clara, CA, USA).

Typical mass spectrometric conditions included gas 1, nitrogen (40 psi); gas 2, nitrogen (40 psi); ion spray voltage, 5,000 V; ion source temperature, 450 °C; curtain gas, nitrogen (25 psi). Multiple reaction monitoring (MRM) scanning in positive ionization mode was used to monitor the transition of m/z 531.201 to 398.191 for NA-184 and 528.149 to 241.100 for NA-101. Gradient elution was carried out consisting of acetonitrile (A) and aqueous buffer (B) (0.1% formic acid containing 2 mM ammonium acetate) with a flow rate of 0.3 ml/min. The elution began at 20% of mobile phase A and maintained for 1.5 min. The ratio was increased to 90% in 0.5 min and kept for 6 min. Afterwards, the gradience was returned to to 20% of A and balanced for 1.5 min. The temperatures of analytical column and autosampler were both set at room temperature.

This method showed a good linearity for concentration ranging from 2 to 500 ng/ml with average accuracy between 85% to 115%. The lower limits of quantification (LLOQ) for plasma samples and tissue samples were set as 2 and 5 ng/ml, respectively.

## Results

### 1. Molecular modeling, synthesized molecules, and screening of novel calpain inhibitors

Using the calpain activity assay, we determined that the S-S diastereoisomer of C2I is the active, as compared to the inactive R-S isomer (Fig. 1a). Using the S-S diastereoisomers, we computed the selectivity of known ketoamide calpain inhibitors between calpain-1 and calpain-2 using free energy perturbation (Fig. 1a) (Chatterjee et al., 2017). The correlation between computed and experimental selectivity provided a validation of our *in silico* protein models (Figure 1a). These protein models were then used to rank aromatic fragments at P1’ position using SILCS docking protocol (see Methods). The SILCS FragMaps and docking scores provided a general structure-activity relationship at P1’ site, in which additional substitution is favored at para-position of phenyl group in calpain-1 P1’ site, but at meta-position in calpain-2 P1’ site; hydrogen bond donor and acceptor were present at ortho-position in calpain-2, but no density was found at ortho-position in calpain-1 P1’ site (Figure 1 b).

Based on these analyses, about 130 analogs of C2I (Table I) were synthesized (Table I), and tested for their selectivity and potency against human calpain-1 and calpain-2. Results from a subset of these compounds are shown in Table II.

**Table I.**
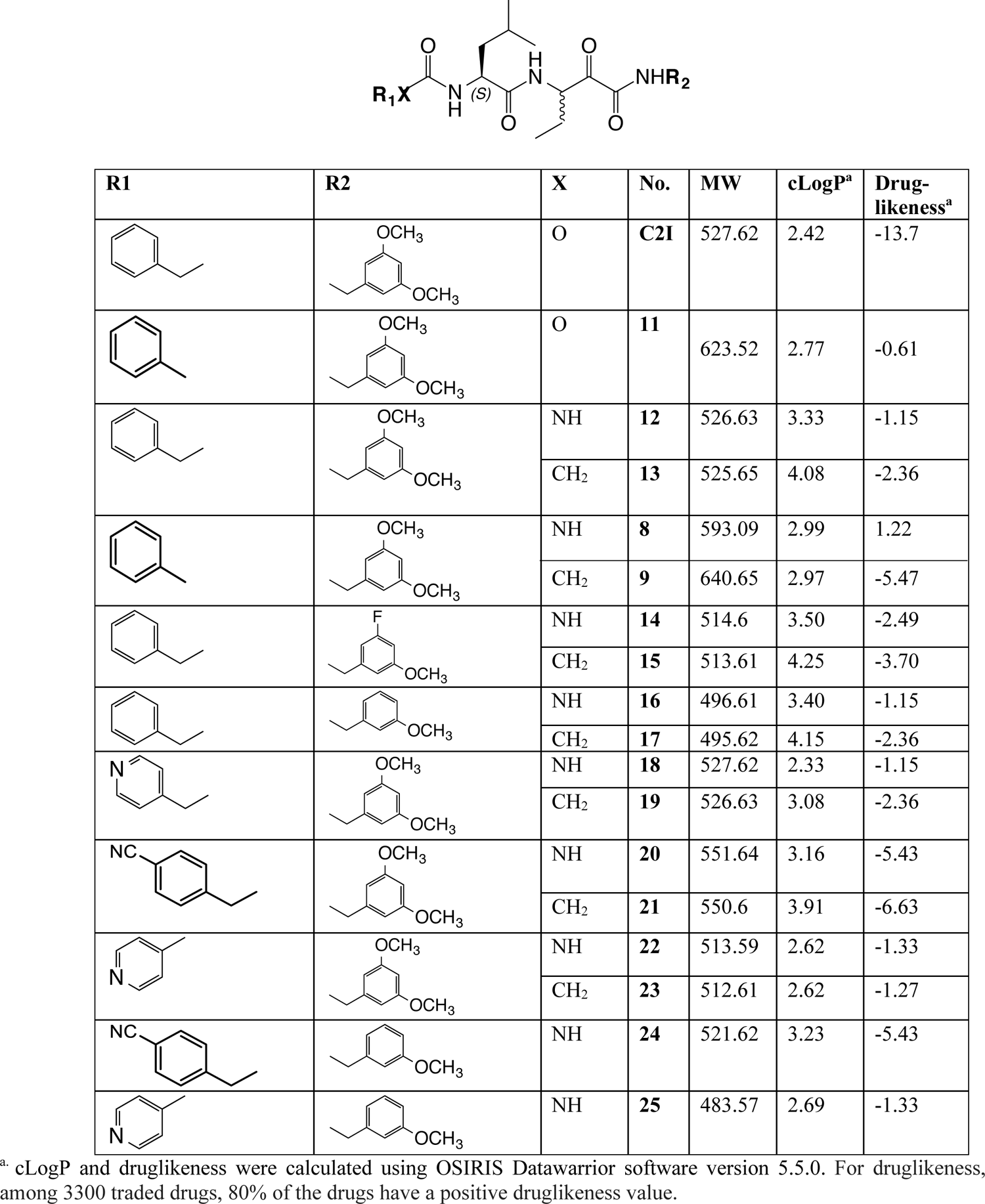
In silico data for various synthesized C21-like compounds.

**Table II:**
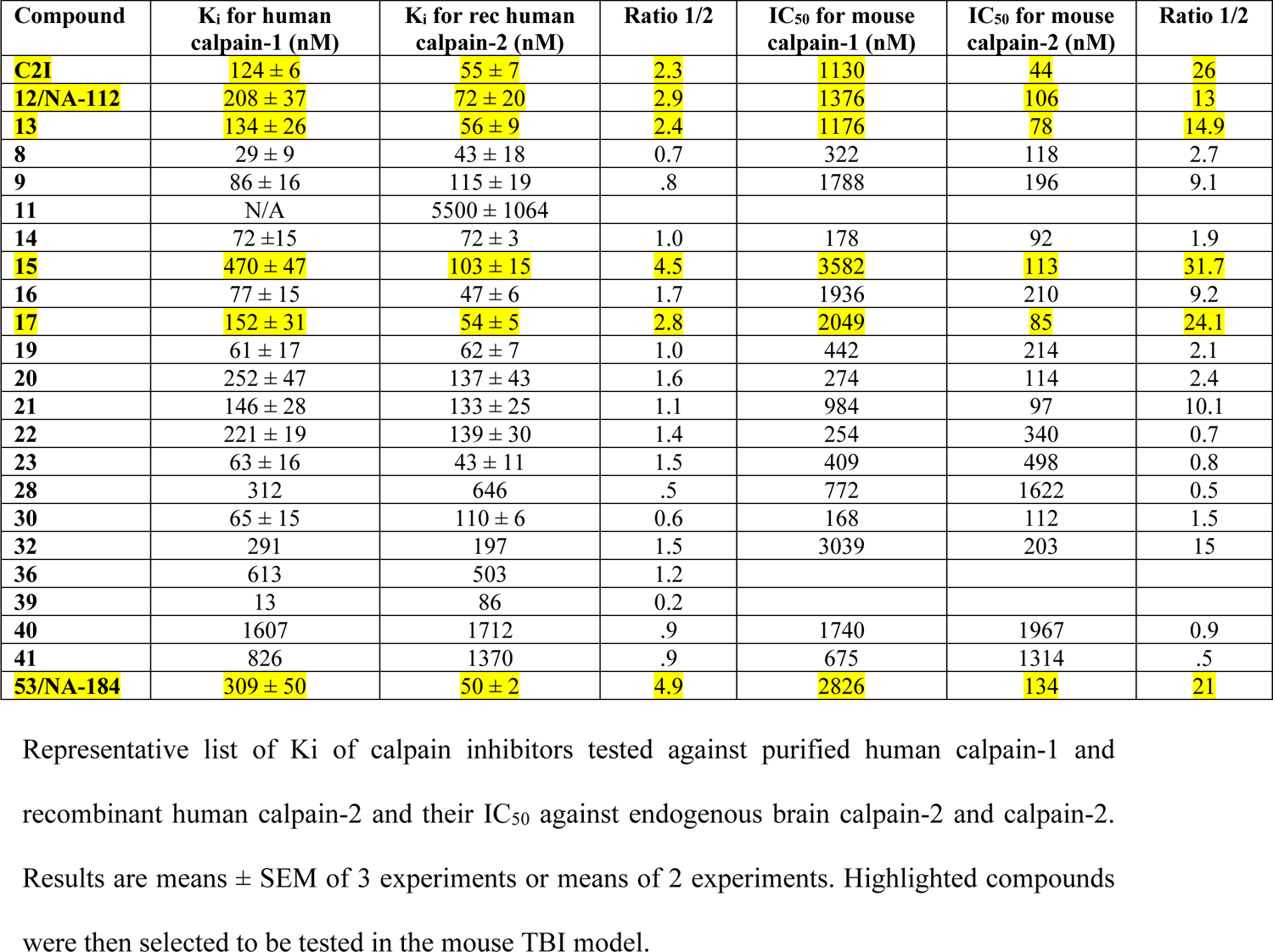
K_i_ and IC_50_ for synthesized compounds.

In addition, the selectivity and potency of these compounds were tested against mouse calpain-1 and calpain-2 activity in cerebellar membrane preparations. Examples for this type of analysis are shown in Fig. 2, where compounds 12 and 53 (aka NA-112 and NA-184) were compared. Interestingly, while the range of values for Ki ratios of calpain-1 over calpain-2 was not wide, the ratios for the IC_50_ values of calpain-1 over calpain-2 showed a much wider range.

**Figure 2:**
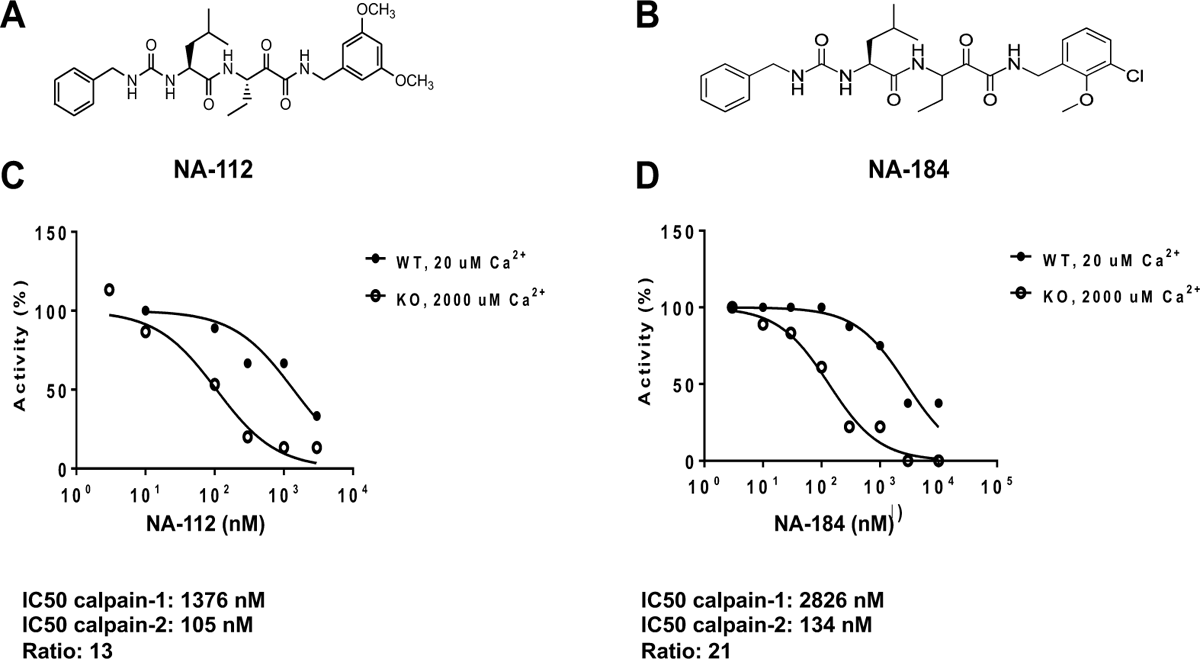
Selectivity of NA-112 and NA-184 for calpain-2 vs calpain-1 A. NA-112 structure. B. NA-184 structure. C. Inhibition of mouse calpain-1 and calpain-2 by NA-112. Membrane fractions from WT or calpain-1 knock-out (KO) mice were prepared. Calpain-1 activity was measured in membrane fractions from WT mice in the presence of 20 µM calcium. Calpain-2 activity was measured from membrane fractions from calpain-1 KO mice in the presence of 2 mM calcium. The experiments were repeated twice with similar results.

We focused further testing on molecules for which the ratio of IC50 was larger than 10, as these molecules were likely to be more selective for calpain-2 than for calpain-1. All these molecules were tested in the CCI mouse model of TBI. Compounds were administered ip 1 h after TBI, the animals were sacrificed 24 h later, then calpain-1 and calpain-2 activities in cerebellar membranes were analyzed. The number of degenerating cells around the lesion site was also analyzed by the TUNEL assay in the same animals. Compounds 13, 15 and 17 were tested first as they all showed a high IC_50_ ratios (Fig. 3). Interestingly, compound 15 had no effect on either brain calpain-1 or calpain-2 or on cell death when administered at 0.1 or 1.0 mg/kg. It also showed no effect on the volume of brain lesion at doses up to 10 mg/kg. In contrast, compounds 13 and 17 did inhibit calpain-2 and cell death at 1.0 mg/kg. Compound 13 appeared to be more potent than compound 17 as it did inhibit brain calpain-2 and cell death at 0.1 mg/kg. Our interpretation for the lack of effect of compound 15 is that this molecule does not cross the blood-brain barrier, as it showed good efficacy in the mouse model of acute glaucoma when injected intraocularly (data not shown).

**Figure 3:**
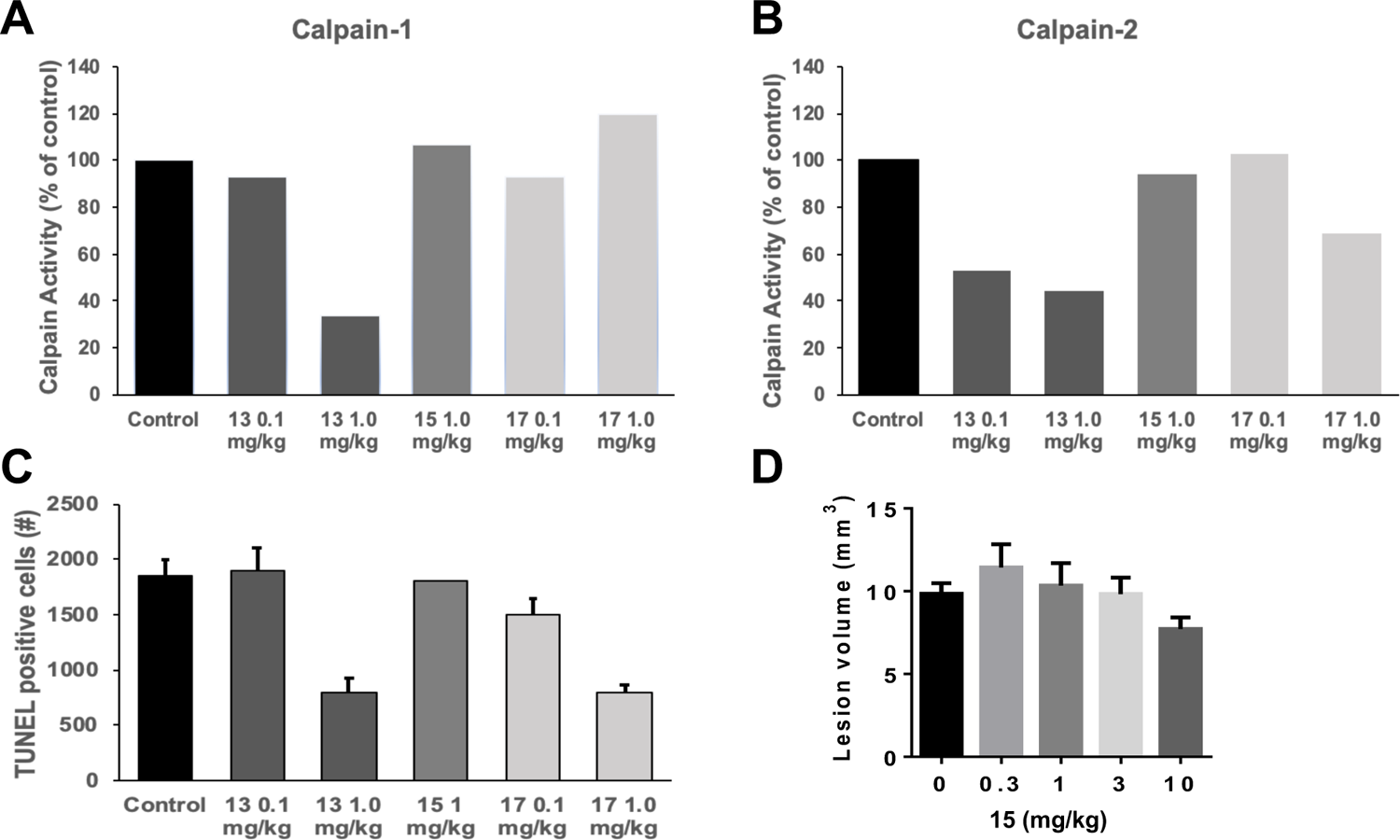
Effects of various calpain inhibitors on calpain activity, cell death and lesion volumes following TBI. Mice were subjected to CCI and were injected 1 h later with the indicated calpain-2 inhibitors (13, 15 and 17) (ip). Twenty-four h later, mice were sacrificed, and calpain activity was measured in cerebellar membrane fractions. Calpain-1 activity represents the activity measured in the presence of 20 µM calcium, while calpain-2 activity represents the difference in activity measured in the presence of 2 mM calcium and 20 µM calcium. The number of TUNEL-positive cells and the lesion volume were determined in tissue sections from the forebrain of the same animals. In A and B results are the means of 2 experiments. In C and D, results are means ± SEM of 3-5 animals.

### 2. Comparison of NA-112 and NA-184 in the mouse model of TBI

Detailed dose-response curves for NA-112 and NA-184 on brain calpain-1 and calpain-2 activity and cell death 24 h after TBI are shown in Figure 4. As previously shown, both NA-112 and NA-184 have higher selectivity for calpain-2 than for calpain-1. The EC_50_ for calpain-2 is 0.11 mg/kg for NA-112 and 0.43 mg/kg for NA-184. NA-112 inhibition of calpain-1 is first detected at 1 mg/kg and is close to 40% at 10 mg/kg. NA-184, however, inhibits less than 10% of calpain-1 activity at 10 mg/kg. The effects of the 2 compounds on cell death are in agreement with their effects on calpain-2 and calpain-1. Thus, low doses of NA-112 prevent cell death at doses up to 1 mg/kg, but higher doses are less effective, in agreement with the inhibition of calpain-1, as previously reported for NA-101 (Wang et al, 2018). In contrast, NA-184 results in a dose-dependent inhibition of TBI-induced cell death with an EC_50_ of 0.13 mg/kg, a maximal effect reached at 1 mg/kg and no further protective effect at doses up to 10 mg/kg. Based on these results, we decided to pursue the pre-clinical development of NA-184 for the treatment of TBI. Note that we have obtained excellent neuroprotective results with NA-112 in a mouse model of epilepsy (Wang et al., 2021), and will therefore consider the potential preclinical development of NA-112 for epilepsy.

**Figure 4:**
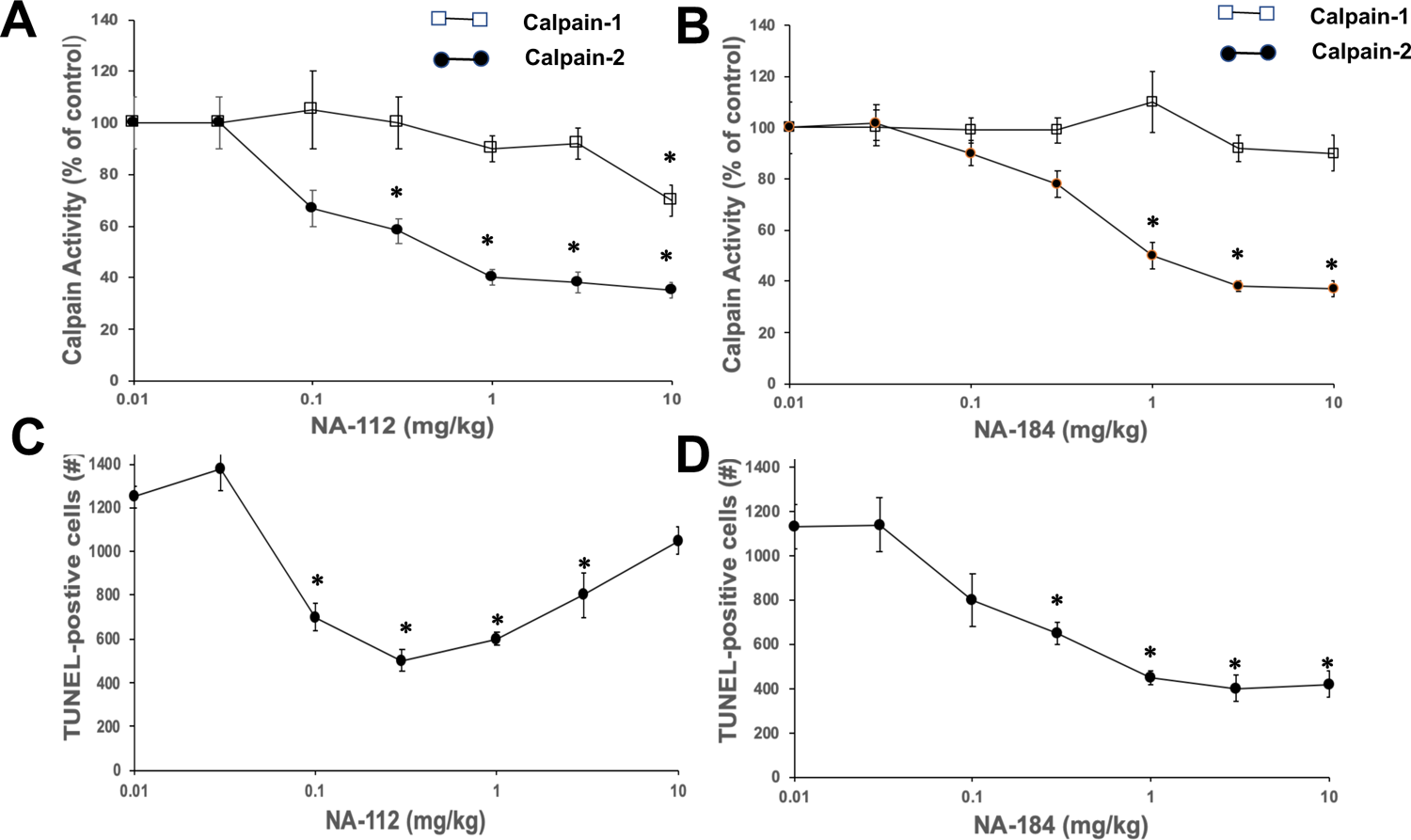
Effects of NA-112 and NA-184 on calpain activity and cell death after CCI. Mice were subjected to CCI and were injected 1 h later with various doses of NA-112 or NA-184 (ip). Twenty-four h later, mice were sacrificed, and calpain activity was measured in cerebellar membrane fractions. Calpain-1 activity represents the activity measured in the presence of 20 µM calcium, while calpain-2 activity represents the difference in activity measured in the presence of 2 mM calcium and 20 µM calcium. The number of TUNEL-positive cells was determined in tissue sections from the forebrain of the same animals. Results are means ± SEM of 4-5 animals.

### 3. Effects of NA-184 in the CCI model of TBI in male and female mice and rats

To further determine the effects of NA-184 as a potential treatment for TBI, we tested its effects in the CCI model of TBI in male and female mice and in male and female rats (Fig. 5). Animals were injected with NA-184 (1 mg/kg, ip) 1 h after TBI and were sacrificed 24 h later. Calpain-1 and calpain-2 activity were assayed in membrane fractions from the cerebellum and cell death in the cortical area surrounding the trauma was analyzed with the TUNEL assay. NA-184 administration had no effect on calpain-1 activity (Fig. 5A) but resulted in 50-70% inhibition of calpain-2 activity in both male and female mice and rats (Fig. 5B). The treatment also resulted in a significant reduction (between 30 to 70%) of cell death in the cortical area surrounding the lesion site (Fig. 5C). The levels of cell death were plotted as a function of the relative activity of calpain-2 in male mice (Fig. 5D); there was a highly significant correlation between the levels of calpain-2 activity and the level of cell death, further supporting the conclusion that calpain-2 activity is responsible for cell death following TBI.

**Figure 5:**
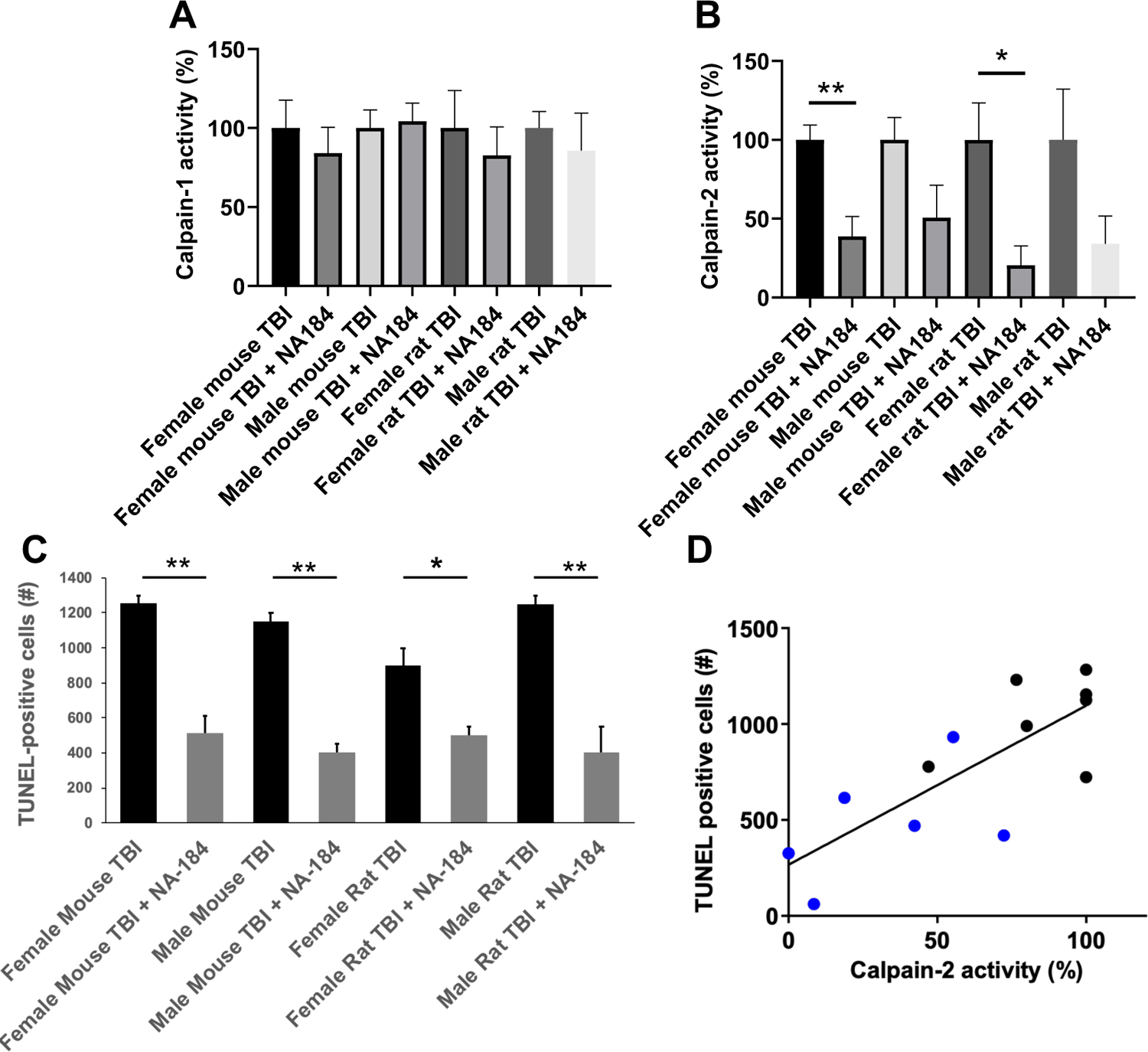
Effects of NA-184 on calpain activity and cell death in male and female rats and mice following TBI. Male and female mice and rats were subjected to CCI and were injected 1 h later with 1 mg/kg NA-184 (ip; rats were injected twice at 1 and 8 h after CCI). Twenty-four h later, mice and rats were sacrificed, and calpain activity was measured in cerebellar membrane fractions. Calpain-1 activity represents the activity measured in the presence of 20 µM calcium (**A**), while calpain-2 activity represents the difference in activity measured in the presence of 2 mM calcium and 20 µM calcium (**B**). The number of TUNEL-positive cells was determined in tissue sections from the forebrain of the same animals (C). D. Correlation between number of dead cells and calpain-2 activity levels in male rats (back dots: CCI; blue dots: CCI + NA-184). Results are means ± SEM of 4-5 animals. * p < 0.05; ** p < 0.01 (Student’s t-test).

### 4. Selectivity, Stability and Epimerization of NA-184

The overall features of NA-184 are reported in Table III. Recombinant human calpain-2 was used in the initial screening of the newly synthesized inhibitors as well as an artificial substrate. The recombinant human calpain-2 includes the full-length human large subunit of calpain-2 but a truncated small subunit. We thought it would be important to determine the inhibitory potency of NA-184 against full-length human calpain-2 and when using an endogenous protein target. Lysates from HEK cells lacking either calpain-1 or calpain-2 were used to test the potency of NA-184 to inhibit the truncation of spectrin. Under these conditions, we found an EC_50_ for NA-184 of 1.55 nM and no inhibition of human calpain-1 at concentrations up to 10 µM (Fig. 6).

**Figure 6:**
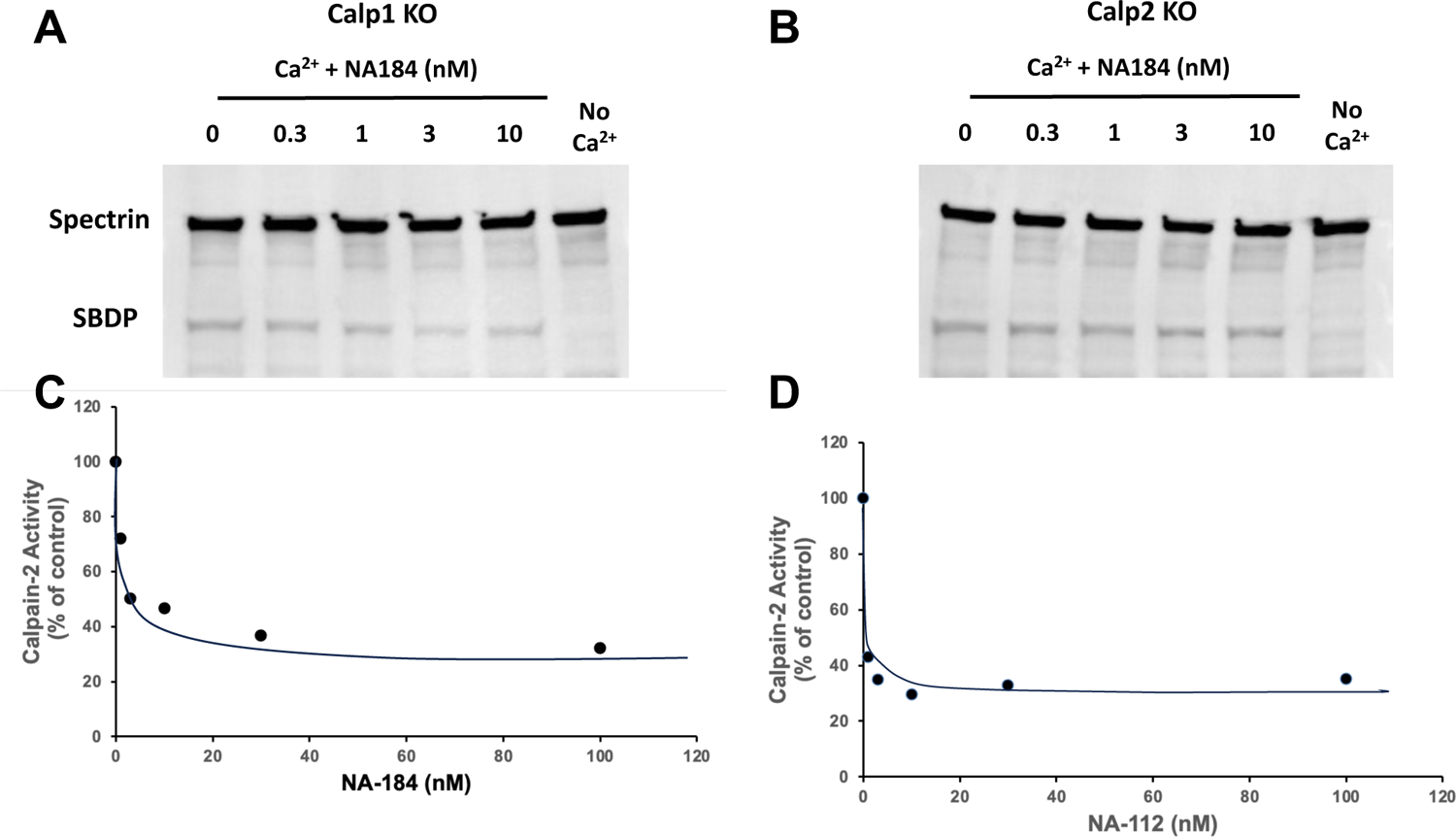
Effects of NA-184 and NA-1112 on human calpain-1 and calpain-2. HEK cells deleted of calpain-1 (Calp1 KO) or calpain-2 (Calp2 KO) were lysed in calpain assay buffer as described in Material and Methods. They were incubated in the presence of 2 mM calcium and the indicated concentrations of NA-184 or NA-112 for 30 min at 30 °C. Aliquots of the homogenates were processed for western blots with a spectrin antibody. Intensities of the native spectrin band and the calpain-mediated breakdown product (SBDP) were quantified and the ratio SPDP/spectrin calculated. Results were normalized to the values found in the absence of calpain-2 inhibitors. The experiment was repeated twice with similar results.

**Table III:** Key properties of NA-184.

|  |  |
| --- | --- |
| Molecular weight | 531 g×mol<sup>-1</sup> |
| cLogP3 | 2.62 |
| Drug-likeness 3 | 2.5692 |
| Solubility in PBS | 14 μg/ml |
| Total Surface Area | 416 |
| Relative PSA | 0.25908 |
| In vivo IC<sub>50</sub> against human calpain-2 | 1.55 nM |
| In vivo Inhibition against human calpain-1 | No inhibition up to 10 μM |
| Eurofin Safety Panel (44 receptors/enzymes) | - CCK1: 97% at 10 μM<br>- Kappa Opiate Receptor: 57% at 10 μM<br>- $\mu$ Opiate Receptor: 58% at 10 μM |

cLogP and drug-likeness were calculated using OSIRIS Datawarrior software version 5.2.1. cLogP value of compound is the calculated logP, which is the logarithm of its predicted partition coefficient between n-octanal and water. The cLogP is calculated as the sum of contributions of every atom based on its atom type. Datawarrior uses 368 atom types and more than 5000 experimental logP values as training set to optimize the contribution values associated with the atom types. The drug-likeness is calculated as the total scores of substructure fragments that are present in the molecule. Datawarrior uses 5300 distinct substructure fragments with associated drug-likeness scores. A positive value states that the molecule contains predominantly fragments which are frequently present in commercial drugs.

The effects of NA-184 in the Eurofins safety panel, which contains 44 receptors/enzymes, were also evaluated. The only significant effects of NA-184 were a 97% inhibition of the CCK1 receptor, a 57% inhibition of the K opiate receptor and a 58% inhibition of the µ opiate receptor, all at 10 µM. Inhibition of various cytochrome P450 enzymes, by 10 µM NA-184 was also tested and resulted in 50 to 65% inhibition of CYP2C9, CYP2C19, and CYP2D6.

The effects of NA-184 on various proteases are shown in Table IV. Only cathepsin L and B were significantly inhibited, with NA-184, Ki’s of 1.9 nM and 220 nM, respectively. NA-184 did not, however, exhibit significant inhibition on a variety of other cysteine or serine proteases at concentrations up to 10 µM.

**Table IV:** Selectivity of NA-184 against other proteases.

| **Protease** | **IC<sub>50</sub>** | **K<sub>i</sub>** |
| --- | --- | --- |
| Cathepsin B | 303 nM | 220 nM |
| Cathepsin L | 2.6 nM | 1.9 nM |
| Caspase-3 | | > 21.1 μM |
| S20 proteasome | | > 10 μM |
| Thrombin | No inhibition up to 10 μM | |
| Kallikrein | No inhibition up to 10 μM | |
| ACE | No inhibition up to 10 μM | |
| MMP-8 | No inhibition up to 10 μM | |

Commercially available kits for the various proteases were used to determine NA-184 inhibition against these various proteases.

**Table V:**
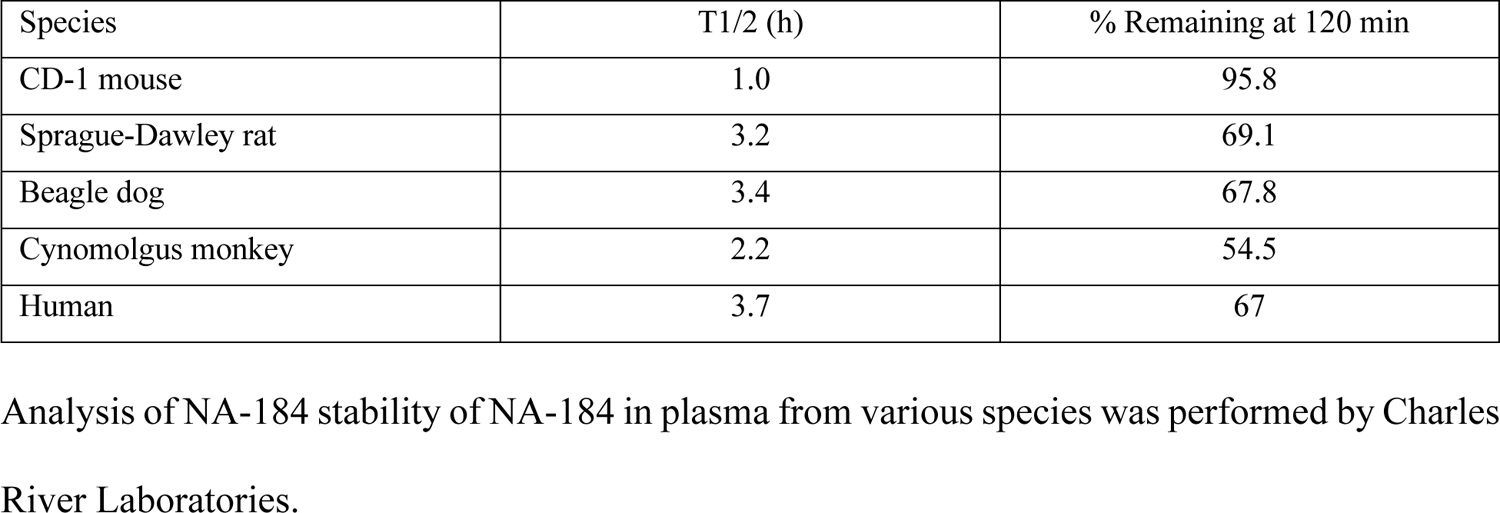
NA-184 stability in plasma from various species.

NA-184 is a mixture of diastereoisomers, as it has 2 chiral centers, the S-S and the R-S isomers. We separated these 2 isomers and determined, using the calpain assay, that the S-S isomer was the active isomer while the R-S isomer was completely inactive. It was therefore important to determine whether the 2 diastereoisomers could epimerize in buffer and/or plasma. The active S-S-diastereoisomer was preincubated in PBS for various periods of time and the remaining inhibitory activity determined. 200 nM was selected as the starting concentration since it represents the Ki of NA-184 against calpain-2. As can be seen in Figure 7A, the inhibitory effect of the active S-S-NA-184 rapidly decreased with a half-life of about 15 min. Conversely, when we preincubated the inactive R-S-NA-184 in PBS and then tested its inhibitory activity against calpain-2, the inactive molecule became active with a half-life of about 15 min (Fig. 7B). Similar effects were found when the 2 compounds were incubated in plasma instead of PBS. Such a rapid epimerization of NA-184 supports the clinical development of the mixture of the two diastereoisomers.

**Figure 7:**
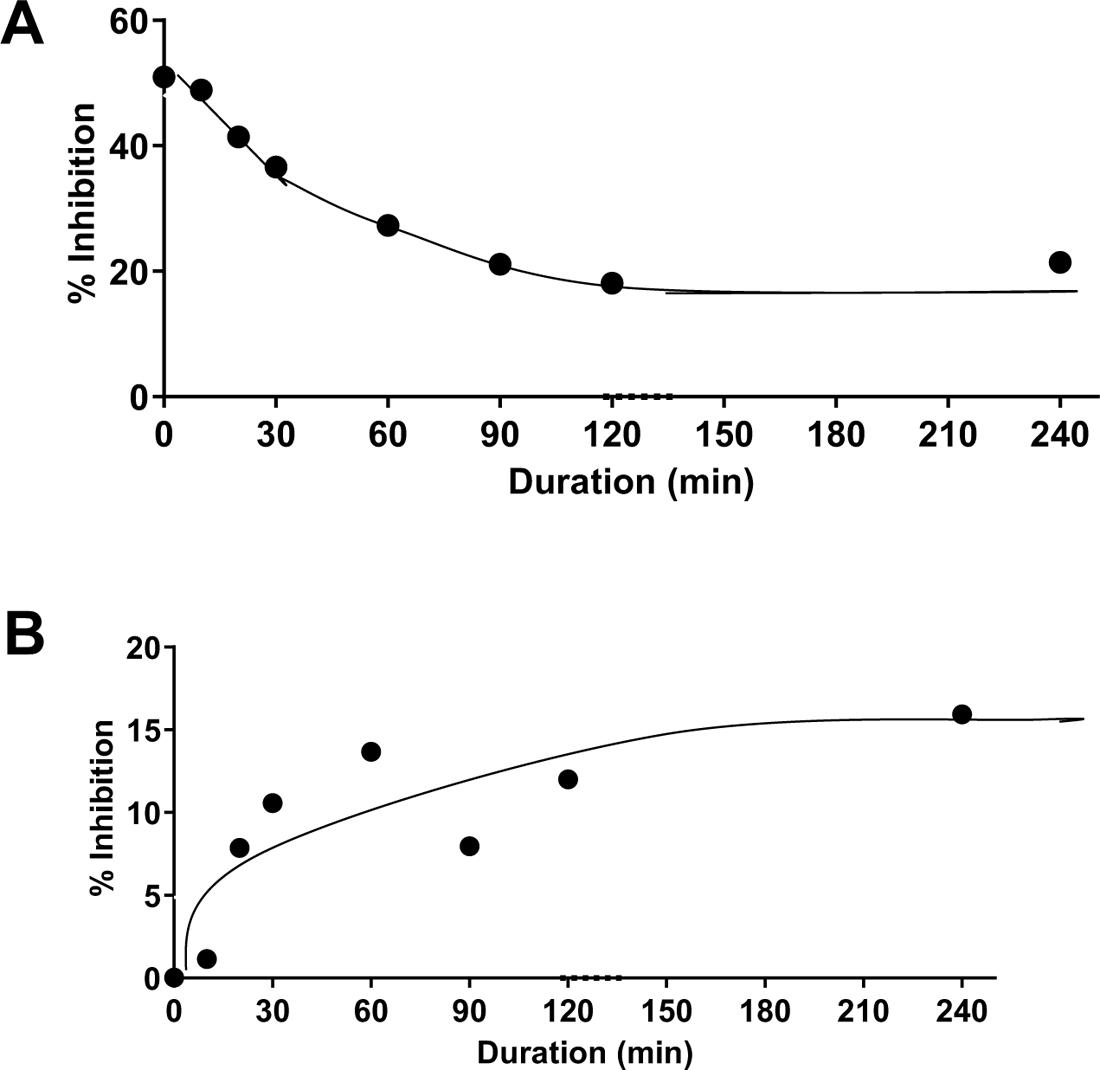
Epimerization of NA-184 in PBS. **A.** The active form of NA-184 (S-S-NA-184, 200 nM) was incubated in PBS at room temperature for the indicated periods of time. Aliquots were then added to the calpain-2 assay buffer and calpain-2 activity determined. **B.** The inactive form of NA-184 (R-R-NA-184, 100 nM) was incubated in PBS at room temperature for the indicated periods of time. Aliquots were then added to the calpain-2 assay buffer and calpain-2 activity determined. The experiment was repeated twice with similar results.

To test the stability in mouse plasma and liver homogenates, NA-184 was incubated in plasma and liver homogenates and aliquots were tested after various periods of time for their ability to inhibit calpain-2 (Fig. 8A,B). The half-life of NA-184 was calculated to be about 3.3 h, while it was about 17.6 h in liver homogenates. These data were extended with studies performed at Charles River Laboratories, which confirmed the relatively short half-life in plasma from different species, including humans, with the exception of the mouse plasma (Table IV).

**Figure 8:**
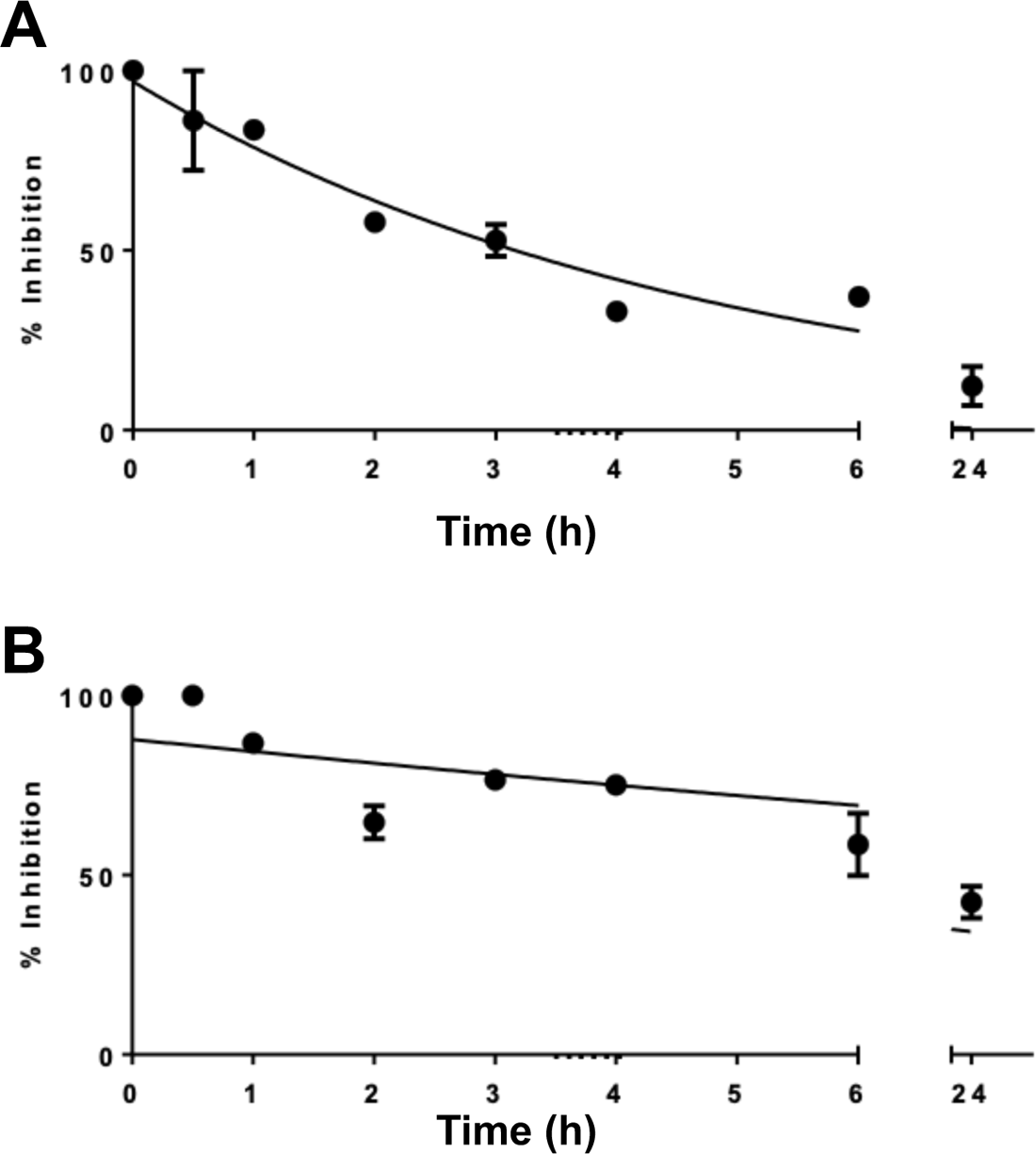
Stability of NA-184 in plasma and liver homogenates. NA184 (0.4 mM) in hydroxypropyl-ß-cyclodextrin (400 mg/ml) was diluted 8 times in freshly prepared mouse plasma (**A**) or freshly prepared mouse liver homogenate (**B**) (final concentration in plasma or liver homogenate: 50 µM). The mixture was incubated at 37 °C for the indicated periods of time. At the indicated time point, 1 µl of the mixture was added to 99 µl of calpain-2 assay solution and calpain-2 activity was determined. As a control, 1 µl of plasma alone was subjected to the calpain-2 assay and its hydrolysis rate was set as 100%. Results are means ± SEM of 3 experiments.

### 5. Pharmacokinetics of N-184 in plasma and brain

We determined the pharmacokinetics properties of NA-184 in C57/Bl mice. The plasma and tissue concentrations versus time curves for both plasma and tissues following iv injection of 10 mg/ml are shown in Fig 9. (Fig. 9). A rapid distribution phase was observed followed by a relatively slow elimination phase with an average elimination half-life of 5.15 h. The typical pharmacokinetic parameters were calculated using a non-compartmental analysis (Table VI). Moreover, NA-184 could rapidly be distributed into brain tissue as the peak concentration was observed at the initial time points. Similarly, a rapid elimination phase in the brain was followed by a slow phase of elimination with a half-life of 3 h (Fig. 9B).

**Figure 9:**
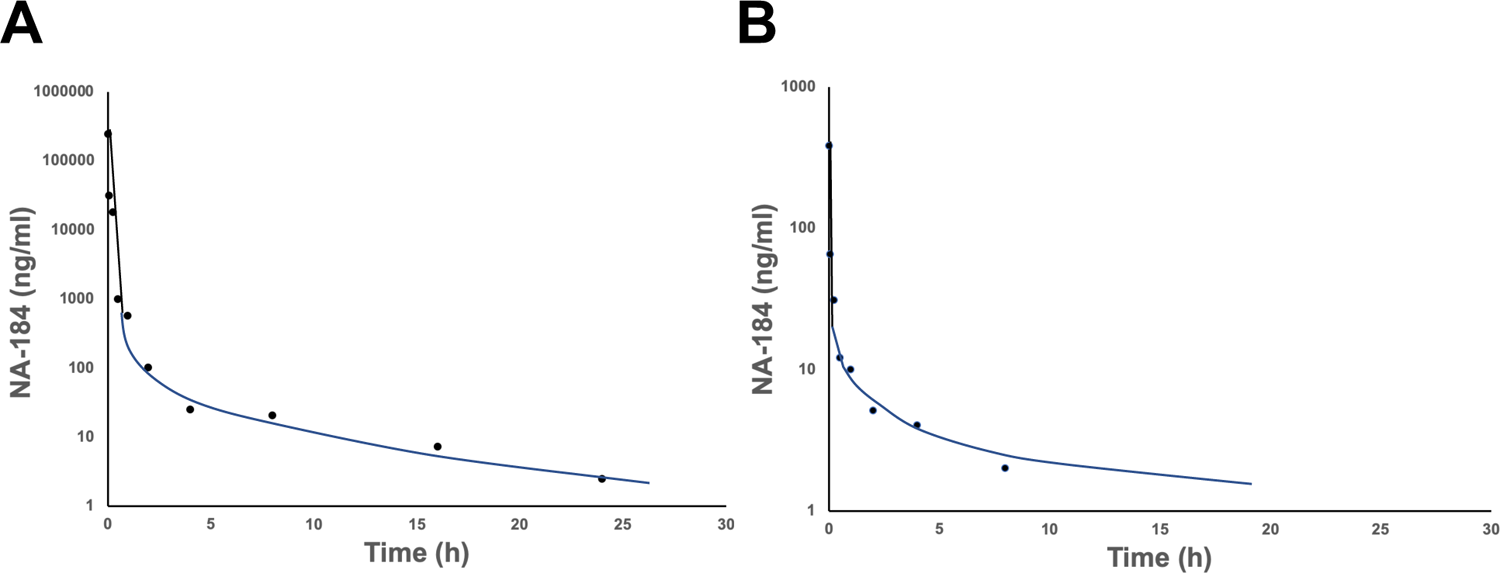
Pharmacokinetics of NA-184 in plasma and brain after iv injection. NA-184 (10 mg/ml in liposomes) was injected in the tail vein of CD1 mice. Mice were sacrificed at various time points, plasma was obtained from retro-orbital blood collection, and whole brains were collected. NA-184 was analyzed as described in Material and Methods. Results are expressed in ng/ml and are the average of duplicate assays.

**Table VI.**
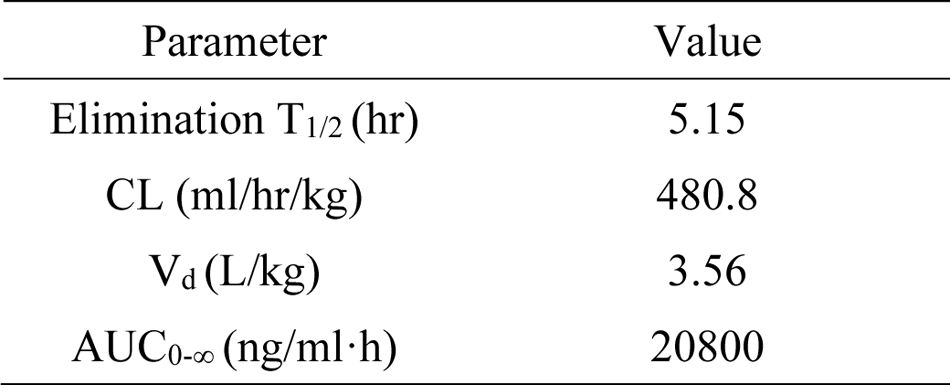
Pharmacokinetic parameters of NA-184 following iv injection (10 mg/kg)

These pharmacokinetic data are useful for understanding the action of NA-184 within the central nervous system, which is essential for therapeutic and safety assessments in the future.

Finally, preliminary studies for in vitro toxicity have been performed and indicated that NA-184 was neither genotoxic nor mutagenic.

## Discussion

Starting from C2I, aka NA-101, we generated about 130 analogs by modifying different aspects of the molecule, mostly on the P1’ and the P3 site. The selectivity of these molecules for calpain-2 versus calpain-1 depended significantly on the source of the enzymes used in the calpain assay. When using purified human calpain-1 and recombinant human calpain-2, the ratios of the Kis against calpain-1 over calpain-2 were not greatly different, although some molecules exhibited 10-fold higher selectivity for calpain-1 and some, 5-fold greater selectivity for calpain-2, thereby resulting in a 50-fold range. The search for better selectivity for calpain-2 than for calpain-1 was facilitated by testing the molecules against mouse brain calpain-1 or calpain-2, which was further enhanced by using the calpain-1 knock-out mouse. This allowed us to identify several molecules with a ratio of the IC50s for calpain-1 over calpain-2 greater than 10. These molecules were then tested in the in vivo mouse model of TBI and led us to the identification of 2 molecules, NA-112 and NA-184, which provided a very significant degree of protection against cell death at relatively low doses, between 0.1 and 1.0 mg/kg. These molecules also exhibited a relatively high degree of selectivity for calpain-2 over calpain-1. In particular, in vivo administration of NA-184 at up to 10 mg/kg did not result in inhibition of calpain-1. NA-184 EC50 against calpain-2 was 0.43 mg/kg and its EC50 for neuroprotection was 0.13 mg/kg; moreover, NA-184 was as effective in male and female rats and mice in the CCI model of TBI. As previously noted (Wang et al., 2018; Wang et al., 2020b), there was a very good correlation between calpain-2 inhibition and protection against neuronal death, further supporting the conclusion that calpain-2 is the calpain isomer responsible for triggering neuronal damage following TBI. As previously reported, deletion of calpain-2 in excitatory neurons of the forebrain also resulted in decreased levels of cell death and lesion volume (Wang et al., 2020a).

While the half-life of NA-184 in plasma was relatively short (about 3 h), it is notable that even 24 h after injection, brain calpain-2 was still significantly inhibited, up to 60-70% following TBI. At this time point, NA-184 levels in the brain would no longer be detectable. However, it is important to stress that these molecules form a reversible covalent bond with the active site of calpain and their in vivo dissociation is probably slow enough that some molecules are still bound to the enzyme at 24 h. Moreover, it is also probably the case that following TBI, the penetration of these molecules across the blood-brain barrier is facilitated (Alluri et al., 2015), resulting in higher brain NA-184 concentrations. Because NA-112 appeared to inhibit calpain-1 at concentrations above 1 mg/kg and loses its neuroprotective effects, we decided to pursue the preclinical development of NA-184 for the treatment of TBI.

NA-184 has several features, which make it a good clinical candidate for TBI treatment. It is very potent against human calpain-2 when tested against spectrin degradation with an EC_50_ of 1.5 nM and does not show inhibition of human calpain-1 at concentration up to 10 µM. Based on our dose-response results in the mouse TBI model where maximal protection was achieved at a dose of 1 mg/kg, we anticipate that this should be dose to be used in the clinic. This converts into a dose of 0.08 mg/kg in human, i.e., 0.15 µM. This concentration is therefore not likely to interfere with the CCK1, kappa and mu opiate receptors, which we found in the Eurofin panel.

NA-184 could potentially interfere with Cathepsin B and L, based on the in vitro assays. At this point we do not know if NA-184 does interfere with these enzymes in vivo, and how it would interfere with the cleavage of their endogenous targets. This issue of selectivity for calpains versus cathepsins has been previously discussed in (Siklos et al., 2015). In this paper, the authors discussed the concept that it is not essential to have a calpain-specific or a cathepsin-specific inhibitor to achieve clinical success in TBI or other neurodegenerative conditions, as both cathepsin and calpain are involved. Interestingly, Cathepsin L inhibitors are being developed for the treatment of various cancers and for Covid. Moreover, Cathepsin L or Cathepsin B knock-out mice do not exhibit any significant neurological problems (Reinheckel et al., 2001). Furthermore, Abbvie recently completed a Phase I clinical trial with their non-selective calpain inhibitor, ABT-957, aka alicapistat, and concluded that twice-daily dosing for 2 weeks did not result in significant adverse effects in both healthy elderly and patients with mild to moderate Alzheimer’s disease (Lon et al., 2019). Finally, it is worth stressing that, for the treatment of TBI patients, relatively short-term treatment (7 days) with NA-184 is planned, which should minimize potential side-effects.

In conclusion, we have identified a potent and selective calpain-2 inhibitor, NA-184, which produces highly significant neuroprotection when administered after TBI in male and female mice and rats. NA-184 has a good drug profile and we are in the process of completing the pre-IND studies in order to initiate clinical trials for the treatment of concussion/traumatic brain injury.

## Acknowledgments

This work was supported by the Office of the Assistant Secretary of Defense for Health Affairs through The Defense Medical Research and Development Program under Award No. W81XWH-19-1-0329. Opinions, interpretations, conclusions, and recommendations are those of the authors and are not necessarily endorsed by the Department of Defense. Grant #BA170606. “Optimization of a selective calpain-2 inhibitor for prolonged field care in Traumatic Brain Injury”. XB is supported in part by funds from the Daljit and Elaine Sarkaria Chair.

## References

Adelman S and Garrison B (1976) Generalized Langevin theory for gas/solid processes: dynamical solid models. Journal of Chemical Physics 65:3751–3761.

Alluri H, Wiggins-Dohlvik K, Davis ML, Huang JH and Tharakan B (2015) Blood-brain barrier dysfunction following traumatic brain injury. Metab Brain Dis 30:1093–1104.

Anagli J, Han Y, Stewart L, Yang D, Movsisyan A, Abounit K and Seyfried D (2009) A novel calpastatin-based inhibitor improves postischemic neurological recovery. Biochem Biophys Res Commun 385:94–99.

Bains M, Cebak JE, Gilmer LK, Barnes CC, Thompson SN, Geddes JW and Hall ED (2013) Pharmacological analysis of the cortical neuronal cytoskeletal protective efficacy of the calpain inhibitor SNJ-1945 in a mouse traumatic brain injury model. J Neurochem 125:125–132.

Baudry M and Bi X (2016) Calpain-1 and Calpain-2: The Yin and Yang of Synaptic Plasticity and Neurodegeneration. Trends Neurosci 39:235–245.

Best RB, Zhu X, Shim J, Lopes PE, Mittal J, Feig M and Mackerell AD, Jr. (2012) Optimization of the additive CHARMM all-atom protein force field targeting improved sampling of the backbone phi, psi and side-chain chi(1) and chi(2) dihedral angles. J Chem Theory Comput 8:3257–3273.

Binder S, Corrigan JD and Langlois JA (2005) The public health approach to traumatic brain injury: an overview of CDC’s research and programs. J Head Trauma Rehabil 20:189–195.

Cagmat EB, Guingab-Cagmat JD, Vakulenko AV, Hayes RL and Anagli J (2015) Potential Use of Calpain Inhibitors as Brain Injury Therapy, in *Brain Neurotrauma: Molecular, Neuropsychological, and Rehabilitation Aspects* (Kobeissy FH ed), Boca Raton (FL).

Chatterjee P, Botello-Smith WM, Zhang H, Qian L, Alsamarah A, Kent D, Lacroix JJ, Baudry M and Luo Y (2017) Can Relative Binding Free Energy Predict Selectivity of Reversible Covalent Inhibitors? J Am Chem Soc 139:17945–17952.

Donkor IO (2011) Calpain inhibitors: a survey of compounds reported in the patent and scientific literature. Expert Opin Ther Pat 21:601–636.

Elce JS, Hegadorn C, Gauthier S, Vince JW and Davies PL (1995) Recombinant calpain II: improved expression systems and production of a C105A active-site mutant for crystallography. Protein Eng 8:843–848.

Gupta R and Sen N (2016) Traumatic brain injury: a risk factor for neurodegenerative diseases. Rev Neurosci 27:93–100.

Hess B, Kutzner C, van der Spoel D and Lindahl E (2008) GROMACS 4: Algorithms for Highly Efficient, Load-Balanced, and Scalable Molecular Simulation. J Chem Theory Comput 4:435–447.

Jo S, Kim T, Iyer VG and Im W (2008) CHARMM-GUI: a web-based graphical user interface for CHARMM. J Comput Chem 29:1859–1865.

Jorgensen W, Chandrasekhar J, Madura J, Impey R and Klein M (1983) Comparison of simple potential functions for simulating liquid water. Journal of Chemical Physics 79:926–935.

Kobeissy FH, Liu MC, Yang Z, Zhang Z, Zheng W, Glushakova O, Mondello S, Anagli J, Hayes RL and Wang KK (2015) Degradation of betaII-Spectrin Protein by Calpain-2 and Caspase-3 Under Neurotoxic and Traumatic Brain Injury Conditions. Mol Neurobiol 52:696–709.

Koumura A, Nonaka Y, Hyakkoku K, Oka T, Shimazawa M, Hozumi I, Inuzuka T and Hara H (2008) A novel calpain inhibitor, ((1S)-1((((1S)-1-benzyl-3-cyclopropylamino-2,3-di-oxopropyl)amino)carbonyl)-3-methylbutyl) carbamic acid 5-methoxy-3-oxapentyl ester, protects neuronal cells from cerebral ischemia-induced damage in mice. Neuroscience 157:309–318.

Li Z, Ortega-Vilain AC, Patil GS, Chu DL, Foreman JE, Eveleth DD and Powers JC (1996) Novel peptidyl alpha-keto amide inhibitors of calpains and other cysteine proteases. J Med Chem 39:4089–4098.

Liu S, Yin F, Zhang J and Qian Y (2014) The role of calpains in traumatic brain injury. Brain Inj 28:133–137.

Liu Y, Wang Y, Zhu G, Sun J, Bi X and Baudry M (2016) A calpain-2 selective inhibitor enhances learning & memory by prolonging ERK activation. Neuropharmacology 105:471–477.

Lon HK, Mendonca N, Goss S, Othman AA, Locke C, Jin Z and Rendenbach-Mueller B (2019) Pharmacokinetics, Safety, Tolerability, and Pharmacodynamics of Alicapistat, a Selective Inhibitor of Human Calpains 1 and 2 for the Treatment of Alzheimer Disease: An Overview of Phase 1 Studies. Clin Pharmacol Drug Dev 8:290–303.

Molecular Operating Environment (MOE) v. 2013.08 (Chemical Computing Group Inc. M, QC, Canada) (2016).

Phillips JC, Braun R, Wang W, Gumbart J, Tajkhorshid E, Villa E, Chipot C, Skeel RD, Kale L and Schulten K (2005) Scalable molecular dynamics with NAMD. J Comput Chem 26:1781–1802.

Reinheckel T, Deussing J, Roth W and Peters C (2001) Towards specific functions of lysosomal cysteine peptidases: phenotypes of mice deficient for cathepsin B or cathepsin L. Biol Chem 382:735–741.

Sasaki T, Kikuchi T, Yumoto N, Yoshimura N and Murachi T (1984) Comparative specificity and kinetic studies on porcine calpain I and calpain II with naturally occurring peptides and synthetic fluorogenic substrates. J Biol Chem 259:12489–12494.

Schoch KM, von Reyn CR, Bian J, Telling GC, Meaney DF and Saatman KE (2013) Brain injury-induced proteolysis is reduced in a novel calpastatin-overexpressing transgenic mouse. J Neurochem 125:909–920.

Siedler DG, Chuah MI, Kirkcaldie MT, Vickers JC and King AE (2014) Diffuse axonal injury in brain trauma: insights from alterations in neurofilaments. Front Cell Neurosci 8:429.

Siklos M, BenAissa M and Thatcher GR (2015) Cysteine proteases as therapeutic targets: does selectivity matter? A systematic review of calpain and cathepsin inhibitors. Acta Pharm Sin B 5:506–519.

Simonson L, Baudry M, Siman R and Lynch G (1985) Regional distribution of soluble calcium activated proteinase activity in neonatal and adult rat brain. Brain Res 327:153–159.

Thompson SN, Carrico KM, Mustafa AG, Bains M and Hall ED (2010) A pharmacological analysis of the neuroprotective efficacy of the brain- and cell-permeable calpain inhibitor MDL-28170 in the mouse controlled cortical impact traumatic brain injury model. J Neurotrauma 27:2233–2243.

Vosler PS, Sun D, Wang S, Gao Y, Kintner DB, Signore AP, Cao G and Chen J (2009) Calcium dysregulation induces apoptosis-inducing factor release: cross-talk between PARP-1- and calpain-signaling pathways. Exp Neurol 218:213–220.

Wang Y, Briz V, Chishti A, Bi X and Baudry M (2013) Distinct roles for mu-calpain and m-calpain in synaptic NMDAR-mediated neuroprotection and extrasynaptic NMDAR-mediated neurodegeneration. J Neurosci 33:18880–18892.

Wang Y, Liu Y, Bi X and Baudry M (2020a) Calpain-1 and Calpain-2 in the Brain: New Evidence for a Critical Role of Calpain-2 in Neuronal Death. Cells 9.

Wang Y, Liu Y, Lopez D, Lee M, Dayal S, Hurtado A, Bi X and Baudry M (2018) Protection against TBI-Induced Neuronal Death with Post-Treatment with a Selective Calpain-2 Inhibitor in Mice. J Neurotrauma 35:105–117.

Wang Y, Liu Y, Nham A, Sherbaf A, Quach D, Yahya E, Ranburger D, Bi X and Baudry M (2020b) Calpain-2 as a therapeutic target in repeated concussion-induced neuropathy and behavioral impairment. Sci Adv 6.

Wang Y, Liu Y, Yahya E, Quach D, Bi X and Baudry M (2021) Calpain-2 activation in mouse hippocampus plays a critical role in seizure-induced neuropathology. Neurobiol Dis 147:105149.

Wang Y, Lopez D, Davey PG, Cameron DJ, Nguyen K, Tran J, Marquez E, Liu Y, Bi X and Baudry M (2016) Calpain-1 and calpain-2 play opposite roles in retinal ganglion cell degeneration induced by retinal ischemia/reperfusion injury. Neurobiol Dis 93:121–128.

Wang Y, Zhu G, Briz V, Hsu YT, Bi X and Baudry M (2014) A molecular brake controls the magnitude of long-term potentiation. Nat Commun 5:3051.

Yildiz-Unal A, Korulu S and Karabay A (2015) Neuroprotective strategies against calpain-mediated neurodegeneration. Neuropsychiatr Dis Treat 11:297–310.

